# CRISPR-Cas9 interference in cassava linked to the evolution of editing-resistant geminiviruses

**DOI:** 10.1101/314542

**Authors:** Devang Mehta, Alessandra Stürchler, Matthias Hirsch-Hoffmann, Wilhelm Gruissem, Hervé Vanderschuren

**Affiliations:** Plant Biotechnology, Institute of Molecular Plant Biology, Department of Biology, ETH Zurich, Universitätstrasse 2, Zurich, 8092, Switzerland; Plant Genetics, AgroBioChem Department, University of Liège, Passage des Déportés 2, Gembloux, 5030, Belgium (current address)

## Abstract

We used CRISPR-Cas9 in the staple food crop cassava with the aim of engineering resistance to *African cassava mosaic virus*, a member of a widespread and important family of plant-pathogenic DNA viruses. We found that between 33 and 48% of edited virus genomes evolved a conserved single-nucleotide mutation that confers resistance to CRISPR-Cas9 cleavage. Our study highlights the potential for virus escape from this technology. Care should be taken to design CRISPR-Cas9 experiments that minimize the risk of virus escape.

---

The bacterial CRISPR-Cas9 (Clustered, regularly interspaced short palindromic repeats-CRISPR associated 9) gene editing system can be used to engineer resistance to DNA viruses through direct cleavage of the virus genome. Unlike conventional gene-editing using CRISPR-Cas9, engineering virus resistance requires constitutive and permanent expression of the ribonucleoprotein complex in the host. For example, CRISPR/Cas9 system has been used to engineer immunity to latent HIV-1 proviruses, Hepatitis B viruses, Herpes simplex virus and the Human papillomavirus in mammalian cell lines^1^. CRISPR-Cas9 has also been used in the model plants *Arabidopsis thaliana* and *Nicotiana benthamiana* to engineer resistance to ssDNA geminiviruses^2–4^. However, the degree to which using CRISPR-Cas9 to engineer virus-resistance results in the evolution of resistant viruses is unknown. One concern (which has previously been highlighted^5^) might be that planting transgenic, virus-resistant CRISPR-Cas9 plants in the field will impose a selection pressure on viruses, while simultaneously providing viruses with a mechanism (via Cas9-induced mutations) to escape resistance. We applied CRISPR-Cas9 to engineer resistance to geminiviruses in cassava and investigated the impact of engineering resistance on geminivirus evolution. We failed to engineer geminivirus resistance, but, we found that use of CRISPR-Cas9 led to emergence of a novel, conserved mutant virus that cannot be cleaved again. We urge caution in the application of CRISPR-Cas9 for virus resistance in plants, both in glasshouse and field settings, to prevent the evolution of novel viruses.

Cassava is a tropical staple food crop consumed by more than a billion people. Cassava production in Africa and South Asia can be decimated by cassava mosaic geminiviruses^6^. Biotechnology has proven effective for engineering cassava mosaic virus resistance by using plant endogenous RNA interference (RNAi) pathways to limit the expression of virus genes^7^. However, the fact that plant viruses have developed effective suppressors of RNAi^8^, as well as the limited success of RNAi-mediated geminivirus resistance suggest that newer methods of engineering resistance are needed. Using orthogonal systems like CRISPR-Cas9 to which plant viruses are unlikely to have developed suppressing mechanisms hence appears particularly attractive.

We designed a set of single-guide RNAs (sgRNAs) using a custom algorithm that combines knowledge about potential off-target effects with published software predicting sgRNA template-cleaving efficiency^9^. We chose sgRNA1, which targets both the viral *AC2* gene coding for the multifunctional TrAP protein involved in gene activation, virus pathogenicity and suppression of gene silencing, and the *AC3* gene coding for the REn protein involved in replication enhancement^10^ (Supplementary Fig. 1 a, b). Selected independent transgenic lines (7 Cas9+sgRNA1, 2 control Cas9-only lines, and wild-type controls (WT)) were tested for transgene expression *in vitro* (Supplementary Fig. 1 c, d) and virus resistance in the greenhouse using an infectious clone of ACMV that was introduced using *Agrobacterium tumefaciens*^7^. No significant differences in disease incidence, symptom severity or virus titres were found between test and control lines (Supplementary Fig. 2 a-c).

**Figure 1:**
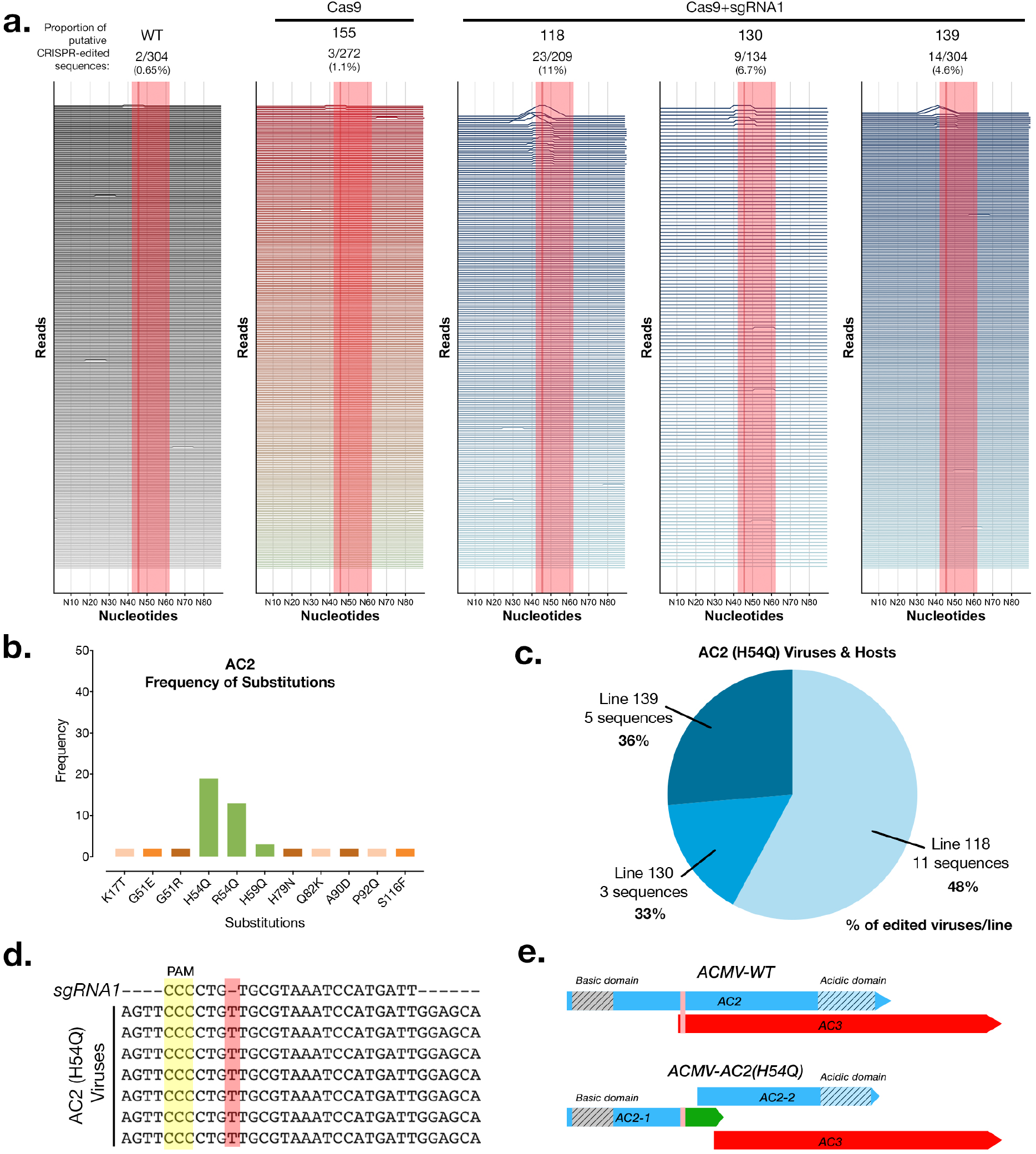
Deep-sequencing of CRISPR-edited viruses in cassava transgenics. **(a)** Analysis of virus sequences from infected plants at 8 weeks post infection. Each horizontal line represents a 90nt window for each individual virus sequence. Peaks represent edits and are scaled to the %mismatch value of each base-pair (see Methods for calculation) in a pairwise global alignment with the reference virus sequence. The sgRNA target is indicated by a shaded red rectangle and a dark line represents the putative cut-site. **(b)** The number of instances of each substitution event in the AC2 protein detected in all the plant lines at 8wpi. Green bars indicate amino acid substitutions in the sgRNA target region. **(c)** Distribution of ACMV-AC2 (H54Q) virus variant in different host plant lines. Percentage values represent the proportion of edited viruses in each line which contain the conserved CRISPR-induced mutation. **(d)** Alignment of some AC2 (H54Q) virus sequences with the sgRNA1 sequence. **(e)** Scheme of the AC2 and AC3 open reading frames in the wild-type and mutant (AC2-H54Q) ACMV virus (vertical pink bar shows the sgRNA target region).

In order to better understand why we couldn’t detect resistance, we sequenced full-length viral amplicons from pooled infected samples at 3 and 8 weeks post-infection (wpi) using Single Molecule Real Time (SMRT) sequencing with a minimum depth of 100 full-length, high quality virus genomes per plant. We chose to sequence the whole viral genome in order to check Cas9+sgRNA1 mediated editing of the targeted viral sequences as well as any other targets. A total of 4942 full-length virus genome sequences were analyzed by alignment against the intact wild-type ACMV genome sequence. We detected CRISPR-edited virus sequences in 3 Cas9+sgRNA1 lines at 8wpi, with line 118 having the highest proportion of edited virus sequences (11%) (Fig. 1 a, Supplementary Fig. 2 d,e, Supplementary Fig. 3). We analysed the AC2 and AC3 genes in depth because these were meant to be edited in Cas9+sgRNA1 lines (Supplementary Fig. 4 & 5). We found that control lines (WT & Cas9), and 5 of 7 Cas9+sgRNA1 lines had similar proportions of substitution [~3%] and indel [~3%] events. In Cas9+sgRNA1 line 118, which had the highest proportion of editing events at 11%, there were more indels [14%] than substitutions [**3%**] (Supplementary Fig. 4). We analyzed the predicted AC2 and AC3 proteins in all sequenced viruses from all of the cassava lines (density plots, Supplementary Fig. 4 & 5). These data indicate that the Cas9+sgRNA1 118 line (and to a lesser extent lines 130 and 139) contain viruses with the desired edits, where the AC2 and AC3 open reading frames stop prematurely at the sgRNA target site. We also detected indels that were present in viruses infecting all plants including controls (minor peaks, Supplementary Fig. 4 & 5, and mismatches, Fig. 1a & Supplementary Fig. 3). These mutations are likely to represent naturally occurring viral variants in our experiment.

While studying the CRISPR-Cas9 caused amino acid substitutions in our entire dataset single-nucleotide insertion in the *AC2* open-reading frame we identified a conserved single-nucleotide insertion in the AC2 open-reading frame (Fig. 1 b-e). Interestingly this conserved mutation was found in three independent Cas9+sgRNA1 lines (but not in any of our 3 control lines). In each of these lines, this virus variant (named *ACMV-AC2 H54Q*) comprised between 33- 48% of all mutant viruses per line, which might indicate selection for this conserved mutation (Fig. 1c). In *ACMV-AC2 H54Q* the single nucleotide ‘T’ insertion (Fig. 1d) results in a H54Q substitution in the AC2 gene and production of a premature stop codon, reducing the length of the AC2 protein from 136AA to 62AA. However, this mutation also creates a new ORF, which might code for the missing AC2 residues. Thus, in this viral variant (Supplementary Data 1), the Cas9-mediated edit at the target site generates two distinct translational units for AC2 (Fig. 1e). Owing to genome organization in cassava geminivirus, the AC3 start codon is disrupted by the *AC2 H54Q mutation*, and the AC3 open reading frame starts at the next ATG, which might result in a truncated 120 AA protein (Fig. 1e). Interestingly the single ‘T’ insertion in *ACMV*-*AC2 H54Q* is in the sgRNA seed sequence. This means that the insertion that is selected for during editing makes *ACMV*-*AC2 H54Q* resistant to further cleavage using sgRNA1, as validated using an *in vitro* cleavage assay (Supplementary Fig. 6).

In our experiment, CRISPR-Cas9 mediated interference of ACMV in cassava transgenic lines resulted in selection for a conserved, abundant, cleavage-resistant mutant virus. We did not observe a clear disease-resistance phenotype associated with the implementation of an CRISPR-Cas9 sgRNA1 virus interference system. Bacteria and archaea use the CRISPR system to defend against viruses and plasmids, but DNA repair mechanisms differ substantially in bacteria and archaea compared with eukaryotes. Most bacteria lack non-homologous end joining (NHEJ) as a DNA-repair mechanism^11^ and cleaved phage/plasmid DNA is usually degraded rather than repaired^12,13^. In eukaryotes NHEJ enables efficient genome editing by effectively repairing cleaved DNA. However, this efficient repair mechanism makes CRISPR-Cas9 mediated virus resistance more prone to evolving mutant viruses. We detected numerous CRISPR-edited viruses whose sequence suggests that dsDNA replicative forms of these viruses have been repaired by NHEJ post-cleavage. Repaired viral genomes have been detected previously in two earlier studies in stable transgenic plants^2,3^, as well as in a study with a viral-vector delivered CRISPR-Cas9 system in model plants^4^ and studies aimed at engineering virus resistance in mammalian systems^14,15^. Although evolution of stable CRISPR-Cas9 cleavage-resistant viruses in plants has been predicted^5^ we now provide confirmation of this in a crop plant. One additional consideration in experiments to engineer virus resistance in eukaryotes is the monitoring period, which in this study was twice as long as in previous studies^2,3^. We first detected *ACMV-AC2 H54Q* at 8wpi, suggesting that evolution of resistant viruses may occur later in the infection process.

We found that using CRISPR-Cas9 with a single guide RNA in cassava resulted in the emergence of a CRISPR-resistant virus strain (with a single conserved mutation) across three independent transgenic lines. We caution that CRISPR-induced virus evolution could have important implications for field studies using CRISPR-Cas9 to engineer plant virus immunity. The ability of CRISPR-systems to trigger the evolution of new viruses would also impact the regulatory mechanisms available for testing and releasing such plants. While the regulation of non-transgenic plants produced through CRISPR-Cas9-editing is being clarified in the US, Canada and the EU, the regulation of transgenic plants constitutively expressing Cas9 has not yet been considered. We expect that such plants will likely proceed under existing biosafety GMO regulations. However, our results point to a novel environmental containment consideration for regulating the release of plants constitutively expressing Cas9 and sgRNAs targeting a virus. We highly recommend sequencing the virus population in on-going studies utilizing CRISPR-Cas9 to engineer virus immunity in plants and animals. We also recommend testing whether using more efficient versions of Cas9, targeting multiple virus genes or using paired Cas9-nickases to delete larger portions of the viral genome can delay the emergence of resistant virus without enhancing recombination frequencies, which can participate in the emergence of hypervirulent geminivirus isolates^6,16^.

Our findings lead us to conclude that strategies to use CRISPR-Cas9 to engineer virus resistance should be optimized to reduce the emergence of editing-resistant viruses. We have not yet tested the ability of the evolved virus to replicate independently. However, this mutant may also be an intermediate step towards the development of a truly pathogenic novel virus. One important feature of the CRISPR system in bacterial cells is its ability to adapt to invading DNA/RNA species. Further investigation into the mechanisms that underlie foreign DNA recognition and spacer acquisition by CRISPR systems in prokaryotes may result in the successful engineering of adaptive immunity to viruses in plants. In the meantime, the implementation of technologies with the potential to speed up virus evolution should be carefully assessed as they pose significant biosafety risks.

## METHODS

### Plasmid Cloning

The GoldenGate/MoClo system^17^ was used to construct binary vectors for plant transformation. Level 1 plasmids bearing 35S promoter driven *hptII* (selectable marker) and plant codon optimised *cas9-GFP* genes were kindly provided by Dr Simon Bull (ETH Zurich). T7 promoter-driven sgRNA cassettes for *in vitro* expression were chemically synthesised (ThermoFisher) and blunt end cloned into a pJET1.2 vector (ThermoFisher). The U6 promoter-driven sgRNA1 gene was chemically synthesised and directly cloned into a GoldenGate Level 1 vector. Level 1 vectors carrying the *hptII*, *cas9-GFP* and *sgRNA1* genes were cloned into a Level 2 vector via a one-pot reaction to produce binary vectors containing the *hptII* and *cas9* expression cassettes as well as vectors containing the *hptII, cas9* and *sgRNA1* expression cassettes.

### sgRNA Design

sgRNAs targeting the ACMV DNA A were designed using two parameters, cleavage efficiency and potential off-targets. Only sgRNAs with at least two seed sequence mismatches against the entire 750 Mb cassava genome were selected. First, high efficiency sgRNAs were obtained using published software from the Broad Institute, MIT^9^. The mismatch search included screening potential sgRNA targets in the cassava reference genome v6.1 (https://phytozome.jgi.doe.gov) with either the canonical NGG and non-canonical NAG protospacer-associated motifs (PAMs) for *SpCas9*. This yielded only 10 sgRNA designs (out of a total of 305 possible sgRNAs) meeting the off-target screening criteria, of which we selected the best six based on efficiency scores and target locations. These six sgRNAs (Supplementary Fig. 1a) were further tested for effectiveness using an *in vitro* cleavage assay with purified Cas9/sgRNA complexes to cleave full-length AMCV amplicons (Supplementary Fig. 1b). We selected sgRNA1 for stable expression in transgenic cassava because it had the best predicted efficiency and performed well in the *in vitro* cleavage assay.

### *In vitro* Transcription

T7::sgRNA expression cassettes were amplified from their respective pJET1.2 host vectors using primers listed in Supplementary Table 1. One μg of gel-purified linear DNA was used as a template in an *in vitro* transcription reaction using 100U of T7 RNA Polymerase (EP0111, ThermoFisher), 0.1U inorganic Pyrophosphatase (EF0221, ThermoFisher), 40U RiboLock RNase Inhibitor (EO0381, ThermoFisher) and 10mM NTP mix (R0481, ThermoFisher) in 1X T7 RNA Polymerase buffer for 16 hours at 37C. Transcribed sgRNAs were purified by Phenol:Chloroform extraction.

### *In vitro* Cas9 cleavage assay

The *in vitro* Cas9 cleavage assay was performed according to a previously published protocol^18^. ACMV templates for cleavage were purified via PCR amplification from total DNA extracts of infected WT plant tissue using primers listed in Supplementary Table 1. The purified GFP-tagged Cas9 endonuclease was kindly provided by Prof Martin Jinek (University of Zurich). Two hundred and fifty nano-grams of ACMV template DNA was digested with 1uM each of purified sgRNA and Cas9 protein. Digestion reactions were stopped at 15, 60 and 105 minutes. The *in vitro* cleavage assay to test resistance to the Cas9-sgRNA1 complex was similarly performed using a 409bp synthetic dsDNA template (Supplementary Data 2) designed from the ACMV-AC2 H54Q sequence.

### Plant transformation and growth conditions

We generated cassava transgenic lines expressing the Cas9 protein together with sgRNA1 (referred to as Cas9+sgRNA1 lines) as well as control lines expressing only the Cas9 protein (referred to as Cas9 lines) using an established *Agrobacterium tumefaciens-*mediated transformation protocol^19^. Twenty transgenic cassava lines were characterized for T-DNA copy number (Supplementary Fig. 7) and expression of the full-length, 180kDa GFP-tagged, plant-codon optimized Cas9 protein was verified via Western blotting (Supplementary Fig. 8). The selected lines expressed both the Cas9 and sgRNA transgenes at varying levels (Supplementary Fig. 1c,d).

Agrobacterium mediated transformation of cassava variety 60444 was performed according to a previously published protocol. *In* vitro transformed cassava plantlets were grown at 28°C in a 16h photoperiod and sub-cultured at 4 week intervals in CBM media (1X Murashige-Skoog medium, 2% sucrose, 2μM copper sulphate, 0.3% gelrite, pH 5.8). 30 day-old plantlets were transferred to soil and grown in glasshouse conditions (14h photoperiod, 60% relative humidity, day/night temperatures: 26°C/17°C).

### Southern Blotting for T-DNA integration analysis

Total DNA was extracted from leaves harvested from 4 week old *in vitro* grown plants using a modified CTAB (cetyl trimethylammonium bromide) protocol^20^. The same leaf samples were used for Western blots and Reverse Transcription-quantitative PCR (RT-qPCR) analysis. Ten μg of total DNA was restriction digested with 20U of HindIII (ThermoFisher) in an overnight reaction. The digested DNA was separated on a 0.8% agarose-TAE gel and transferred overnight onto a positively charged nylon membrane (Roti-Nylon Plus, Carl Roth). A 700bp probe against the *hptII* gene was PCR amplified from the binary vector and labelled with [α-32P] dCTP using the Prime-A-Gene kit (Promega). The nylon membrane was treated with PerfectHyb Plus Hybridization Buffer (Sigma-Aldrich) for 30 minutes followed by hybridisation together with the radio-labelled probe. Blots were developed using a Typhoon FLA 7500 imaging system (GE Healthcare Life Sciences).

### Western Blotting

Crude protein extracts were prepared by incubating ground leaf tissue in a 5%SDS, 125mM Tris-HCL (pH6.8), 15%glycerol buffer with 1X EDTA-free Complete Protease Inhibitor (Roche) for 20 minutes at room temperature. Samples were centrifuged at 4C for 10 minutes to clear debris. Fifty μg of total protein extract were electrophoresed (after a 1:1 dilution with a bromophenol blue solution) on a pre-cast Novex 4-20% Tris-Glycine Midi gel (ThermoFisher) and transferred to a PVDF membrane (Carl Roth) using a TransBlot Cell (Bio-Rad) system according to the manufacturer’s instructions. Membranes were blocked using a 5%milk and 1XTris buffered saline, 0.1%Tween20 (1XTBS-T) solution for 1 hour at room temperature. The blocked membrane was incubated in a primary blotting solution (1XTBS-T) with SpCas9 monoclonal (mouse) antibody (Diagenode) at a 1:2500 dilution and a polyclonal Actin (Rabbit) antibody (Agrisera) at a 1:1000 dilution for 1 hour at room temperature. After three 5 minute washes with 1XTBS-T, the membrane was incubated with a secondary blotting solution containing IRDye 800CW Goat anti-Mouse IgG and IRDye 680RD Goat anti-Rabbit IgG antibodies at a 1:5000 dilution each. Blots were imaged using the LICOR Odyssey CLx fluorescence imaging system. A PageRuler Prestained protein ladder (ThermoFisher) was used for size estimation.

### Reverse Transcription-quantitative PCR

1.5ug of total RNA extract from leaf samples were DNase 1 treated and reverse transcribed with random hexamer primers using the Revert-Aid First strand cDNA synthesis kit (ThermoFisher) according to the manufacturer’s instructions. Quantitative PCR was carried out with the Fast SYBR-Green dye for 40 cycles on a Lightcycler 480 instrument (Roche). Relative quantitation was performed using the *MePP2A* reference gene using the primers listed in Supplementary Table 1. Results of transgene-expression quantitation using RT-qPCR are presented in Supplementary Fig. 1 c, d)

### Virus inoculation and symptom monitoring

*Agrobacterium tumefaciens* (strain LB4404) cells carrying infectious clones of ACMV-NOg DNA A and DNA B^7^ were cultured for 48h at 28°C in 5 mL YEB broth (5 g/L tryptone, 1 g/L yeast extract, 5 g/L nutrient broth, 5 g/L sucrose, 2mM MgSO4) supplemented with antibiotics (25 mg/L rifampicin, 100 mg/L carbenicilin and 50 mg/L kanamycin). Two ml of the starter cultures were then individually added to 200 mL YEB medium with antibiotics and incubated overnight at 28°C to an OD600 of 0.6-1. Cells were pelleted by centrifuging at 5,000 x g for 10 min, then re-suspended in 5 mL inoculation medium (10 mM MES pH 5.6, 10 mM MgCl_2_, 0.25 mM acetosyringone) and incubated for 2 hours at room temperature with shaking. DNA A and DNA B cultures at an OD_600_ of 2.0 were mixed in equal proportions prior to inoculation.

For inoculation, all leaves save the top leaf were removed from 6-week old cassava plantlets (65 plants in total). The stem and axillary buds were pricked prior to dipping the plantlets in Agrobacterium solution for 10 seconds and subsequently covered in a Plexiglas box for three days. A minimum of 5 inoculated plants per line were monitored for symptom incidence and severity over a period of 8 weeks. Symptoms in the top 3 leaves were scored weekly from 3 to 8- weeks post inoculation (wpi) on a scale of 0-3 as depicted in Supplementary Fig. 9. The first emerging leaf from each plant was harvested 3 wpi. The top three leaves were harvested from each plant at 8 wpi.

### Quantitation of virus titres

Virus quantitation was performed by relative quantification qPCR on 10ng of total DNA extracts using ACMV DNA A specific primers and *MePP2A* genomic DNA reference primers as listed in Supplementary Table 1. Symptomatic leaves were harvested in three separate pools for each plant line (in the absence of symptomatic leaves, asymptomatic leaves were used). Three technical replicates were used per pooled sample. Results are shown in Supplementary Fig. 2c (8 wpi) and Supplementary Fig. 10 (3wpi).

### Single Molecule Real Time sequencing of viral amplicons

Full length viral amplicons from selected plant lines were prepared using target specific primers tailed with universal sequences (Supplementary Table 1) according to the protocol provided by Pacific Biosciences Inc. Equal amounts of each amplicon was used as a template in a second PCR using the Barcoded Universal Primers provided by Pacific Biosciences Inc. (Barcodes used per sample are listed in Supplementary Table 2). The standard SMRTBell library construction protocol was used to prepare a pooled, barcoded, sequencing library. Sequencing was performed using a standard MagBead loading protocol on a PacBio RSII instrument. Polymerase reads were processed into barcode separated subreads by primary analysis on the instrument. The resulting subreads were processed using a standard ReadsOfInsert (ROI) analysis using SMRTPipe with a configuration file specifying a minimum ROI predicted quality of 99.9 and a minimum ROI length of 2600bp.

### Sequence analysis

Sequences representing a near full-length region (2692bp) of the ACMV DNA A genome as well as a 100bp region surrounding the sgRNA target site were extracted from each ROI in order to maintain identical start and end sequence positions in each viral amplicon sequence read. Each resulting sequence was pairwise aligned (global alignment using NEEDLE parameters) against its corresponding reference ACMV-NOg DNA A sequence (GenBank: AJ427910). Pairwise alignment scores were assigned to each nucleotide as the sequence mismatch percentage of a 10- nucleotide window surrounding it. The resulting per-base score (y-axis) along each sequence read was plotted using the ggplot and ggjoy packages in R to produce Fig. 1a and Supplementary Fig. 3. Total pairwise identity scores were used to create Supplementary Fig. 2d,e. Background mismatches resulting from either sequencing errors or viral variants were found in all lines, including controls. We also failed to find any conserved edits on viruses infecting Cas9+sgRNA1 lines (and not control lines), indicating the absence of an off-target on the virus genome (Supplementary Fig. 11). This was expected because the seed sequence of sgRNA1 does not have a close match to another site on the viral genome.

### Code Availability

Python and R scripts used for sequence handling and graphical plotting are freely available at: http://www.github.com/hirschmm/CRISPR_amplicon-seq

### Data Availability

Raw sequence data as well as alignment result files are available at: www.dx.doi.org/10.5281/zenodo.896915. The other raw datasets generated and/or analysed during the current study are available from the corresponding author on reasonable request.

## AUTHOR CONTRIBUTIONS

Conceptualization: DM & HV; Methodology, Investigation and Formal Analysis: DM & AS; Software: MHH & DM; Visualisation, Writing-original draft: DM; Funding acquisition, Resources, Data-analysis, Writing-review & editing, Supervision: HV & WG. All authors agree with the final version of the manuscript.

## ACKNOWLEDGEMENTS

We thank Irene Zurkirchen for professional care of cassava plants in the glasshouse, Andrea Patrignani and the Functional Genomics Center Zurich for assistance with SMRT sequencing, Simon Bull for providing source plasmid DNA, Sukalp Muzumdar for assistance with guide RNA design, Martin Jinek and Caroline Anders for providing pure Cas9-GFP protein, detailed protocols and for helpful discussions. This work was supported by the European Union’s Seventh Framework Programme for research, technological development, and demonstration (EU GA-2013-608422-IDP BRIDGES) and by ETH Zurich.

## Supplementary Information

**Supplementary Figure 1:**
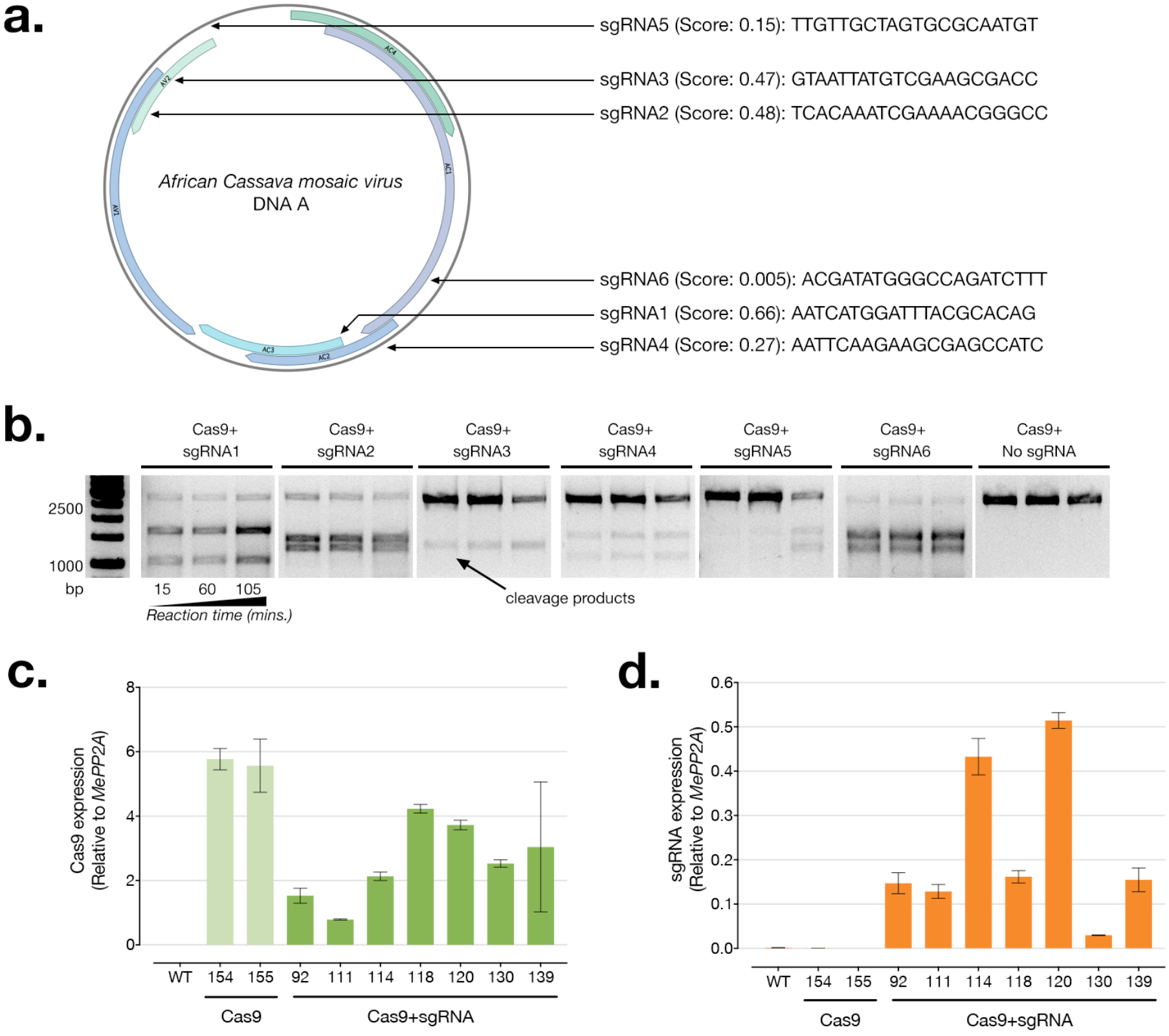
sgRNA design and expression profiling of CRISPR/Cas9 transgenics. **(a)** Low off-target sgRNAs targeting the DNA A of the *African Cassava mosaic virus* and *in silico* predicted efficiency score (max. efficiency=1). **(b)** *In vitro* cleavage assay for testing the effectiveness of six different sgRNAs against the viral template. **(c)** & **(d)** Reverse Transcription quantitative PCR (RT-qPCR) analysis of Cas9 and sgRNA transgene expression respectively. Three independent plants per line were tested.

**Supplementary Figure 2:**
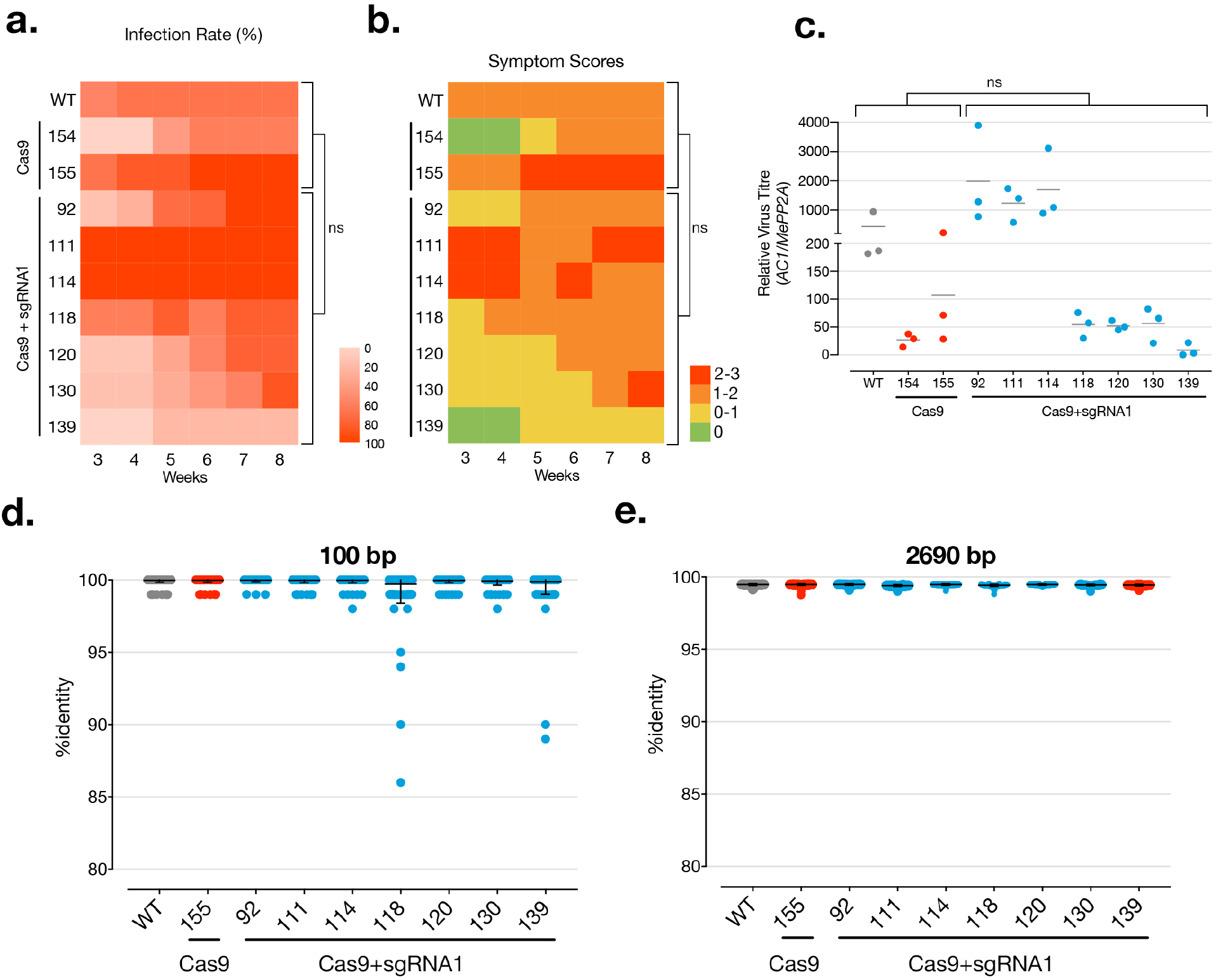
Evaluation of CRISPR/Cas9 expressing transgenics for geminivirus resistance. **(a)** Infection rates of cassava mosaic disease symptoms on agro-inoculated plants monitored weekly over an 8-week period. A minimum of 5 independent replicates per transgenic lines were infected. **(b)** Severity of disease symptoms on agro-inoculated plants. **(c)** Virus levels in symptomatic plant samples at 8 weeks post-infection. Levels were measured on 3 replicated pools of symptomatic leaves from a minimum of 5 individual plants per line. (ns= No statistical significance observed using Dunn’s multiple comparisons test.) **(d)** & **(e)** %identity of each virus sequence per line, measured against the reference sequence, across a 90nt window surrounding the sgRNA site and across almost the entire viral genome respectively.

**Supplementary Figure 3:**
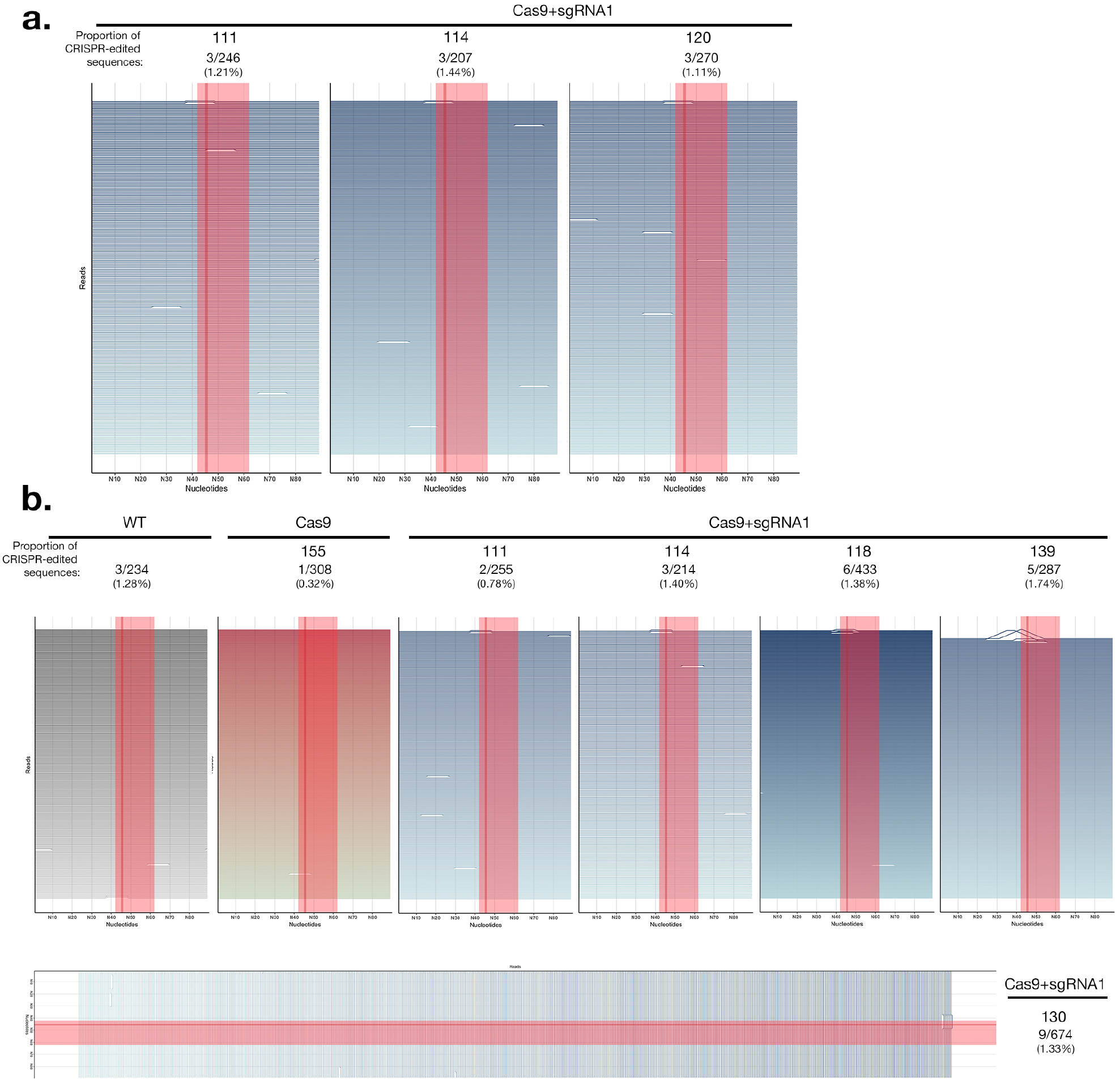
Analysis of virus sequences from infected plants at **(a)** 8 and **(b)** 3 weeks post infection. Each horizontal line represents a 100bp window for each individual virus sequence. Peaks represent edits and are scaled to the %mismatch value of each base-pair (see Methods for calculation) in a pairwise global alignment with the reference virus sequence. The sgRNA target is indicated by a shaded red rectangle and a dark line represents the putative cut-site.

**Supplementary Figure 4:**
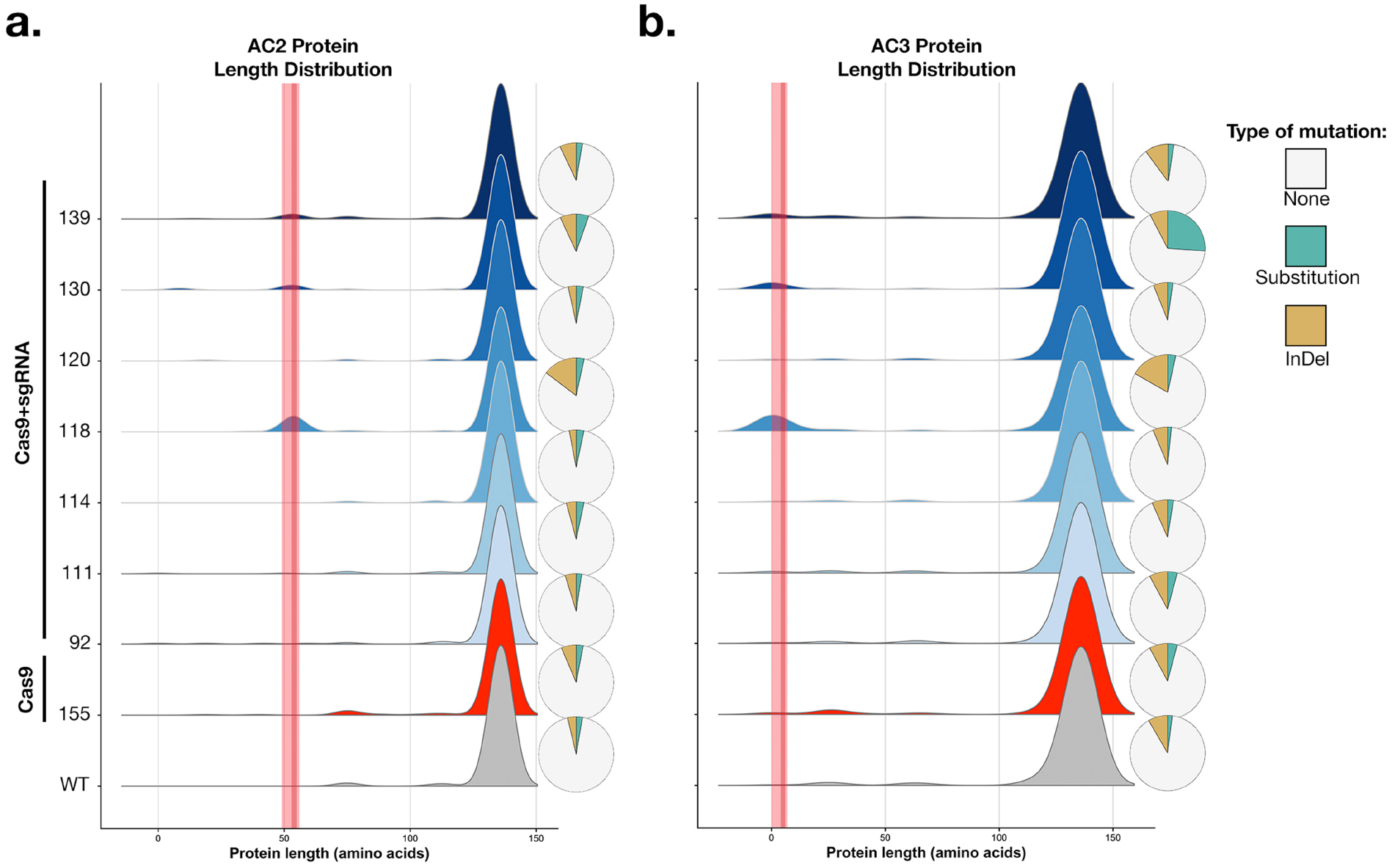
Analysis of viral proteins from edited and control populations obtained by single molecule amplicon sequencing at 8 weeks post infection. **(a)** Density plots show the frequency distribution of virus sequences with different AC2 protein lengths. Pie charts show the type of mutations detected as a percentage of total virus sequences detected in each line. Thus, minor peaks in the distribution show the reduction in protein length caused by the indels (yellow) detected in viruses infecting each line. Red bars show the location of the sgRNA target (light red) and the predicted cleavage site (dark red). **(b)** the same for the AC3 protein at 8 wpi.

**Supplementary Figure 5:**
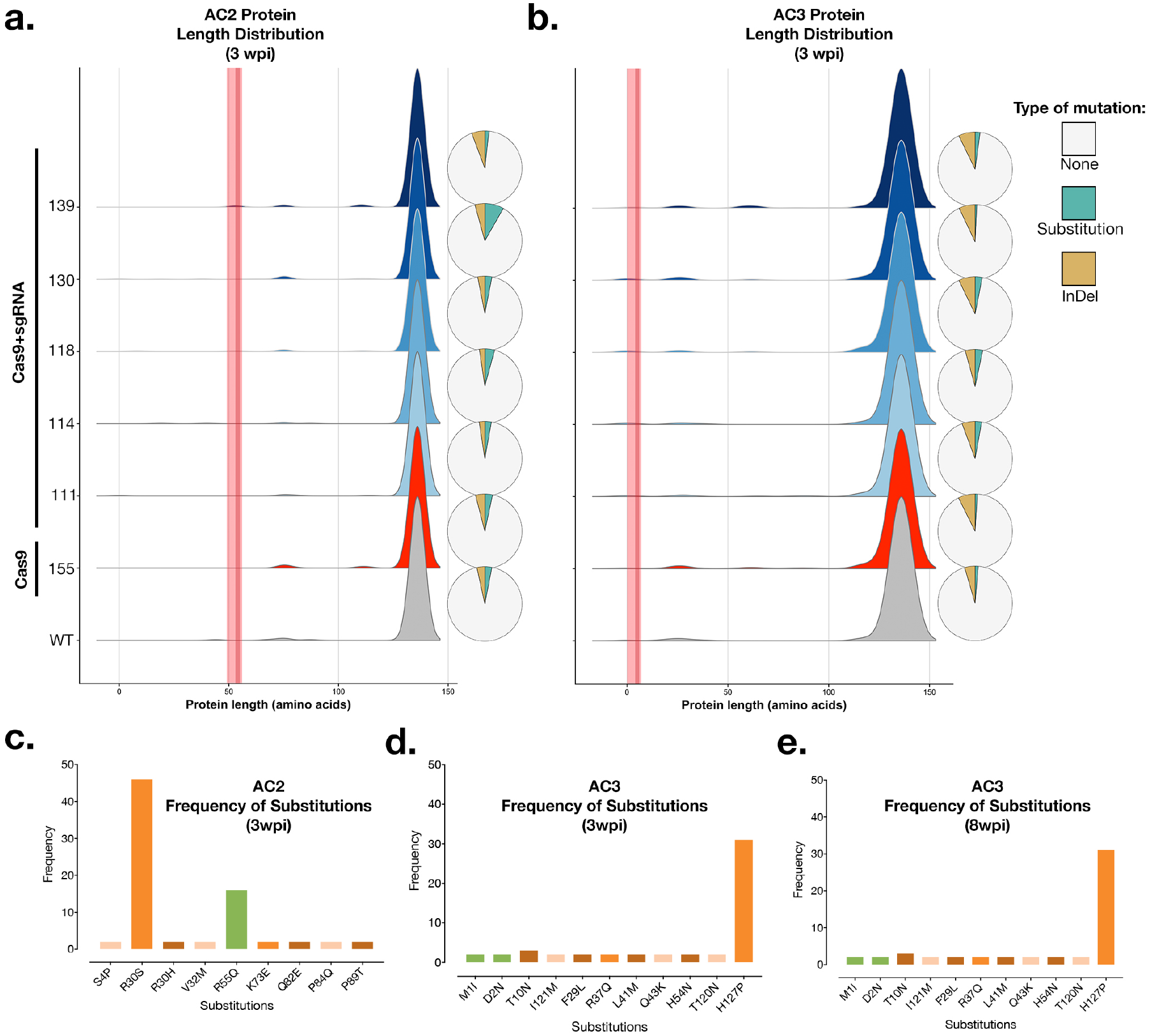
Analysis of viral proteins from edited and control populations obtained by single molecule amplicon sequencing at 3 weeks post infection. **(a)** Density plots show the frequency distribution of virus sequences with different AC2 protein lengths. Pie charts show the type of mutations detected as a percentage of total virus sequences detected in each line. Thus, minor peaks in the distribution show the reduction in protein length caused by the indels (yellow) detected in viruses infecting each line. Red bars show the location of the sgRNA target (light red) and the predicted cleavage site (dark red). **(b)** The number of instances of each substitution event in the AC2 protein detected in all the plant lines. Green bars indicate mutations in the sgRNA target region. **(c)** & **(d)** the same for the AC3 protein at 3 wpi, and **(e)** at 8 wpi.

**Supplementary Figure 6:**
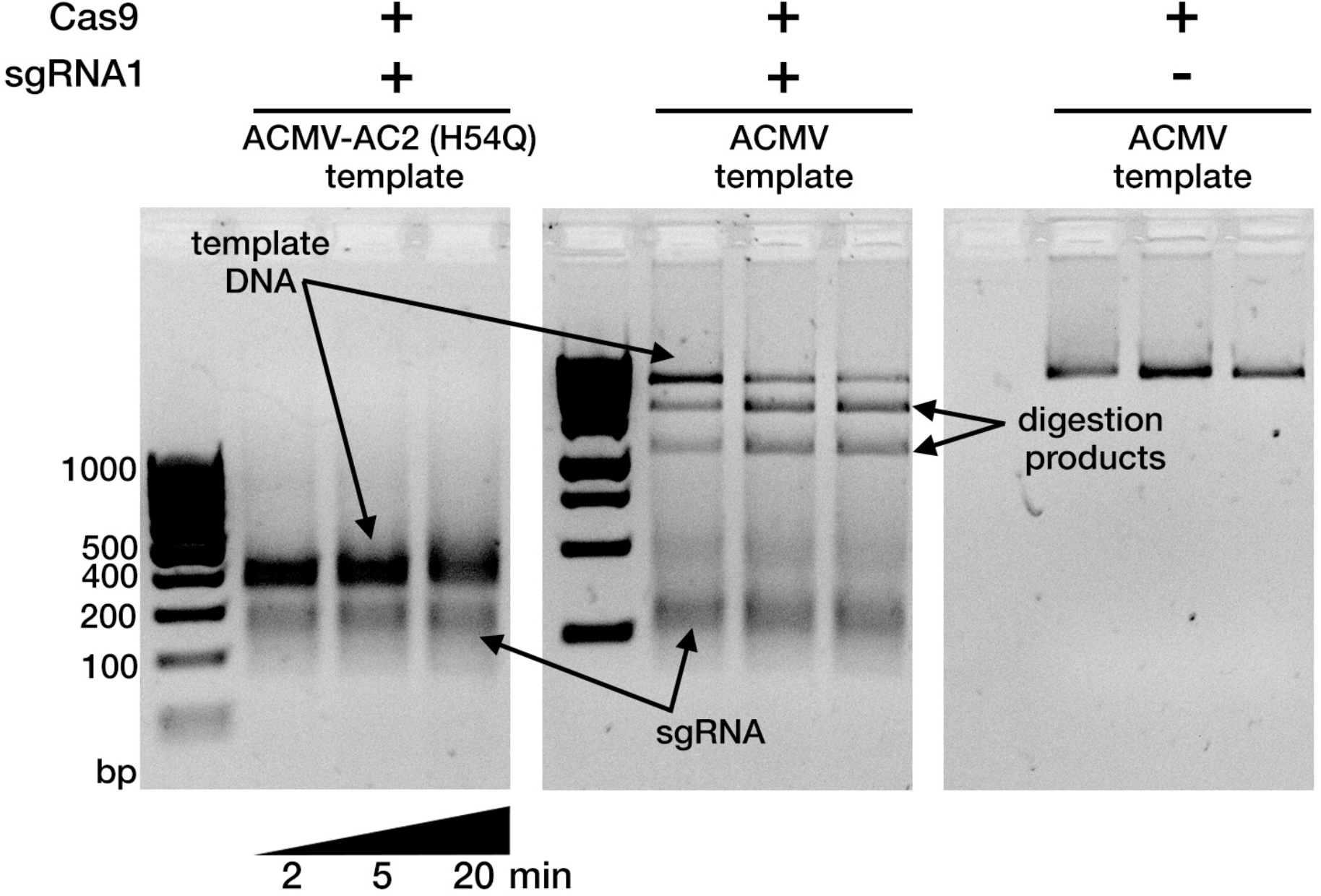
*In vitro* cleavage assay of the ACMV-AC2 H54Q mutant. Treatment of a 409 bp ACMV-AC2 (H54Q) dsDNA template with purified Cas9-GFP and sgRNA1 fails to cleave the ACMV-AC2 (H54Q) amplicon (the expected cleavage products are 248 and 161bp long). The full-length ACMV template used in Fig. 1b is used as positive control and its Cas9-sgRNA1 digestion generates two fragments that are 1696 bp and 1074 bp long.

**Supplementary Figure 7:**
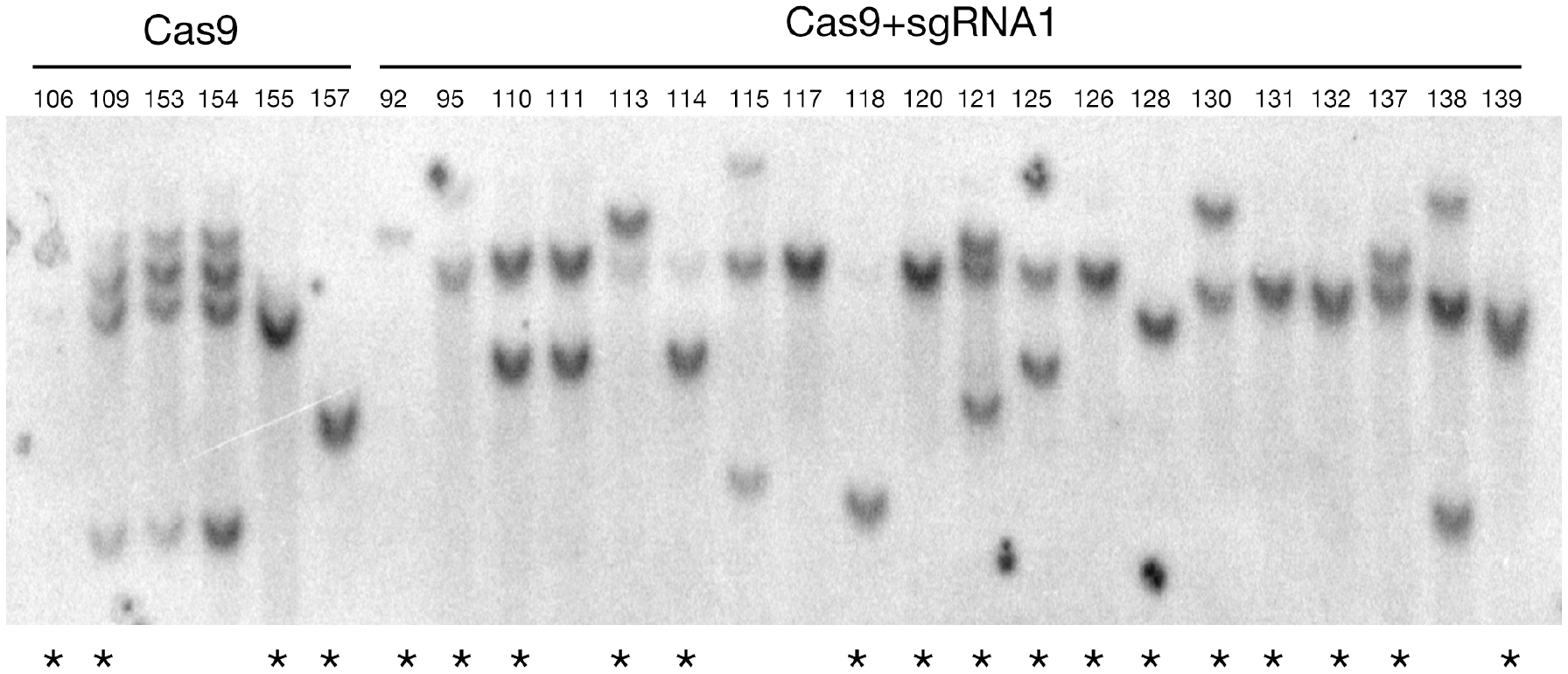
Southern blot analysis for number of T-DNA integration events per plant line. ^*^ indicates independent lines.

**Supplementary Figure 8:**
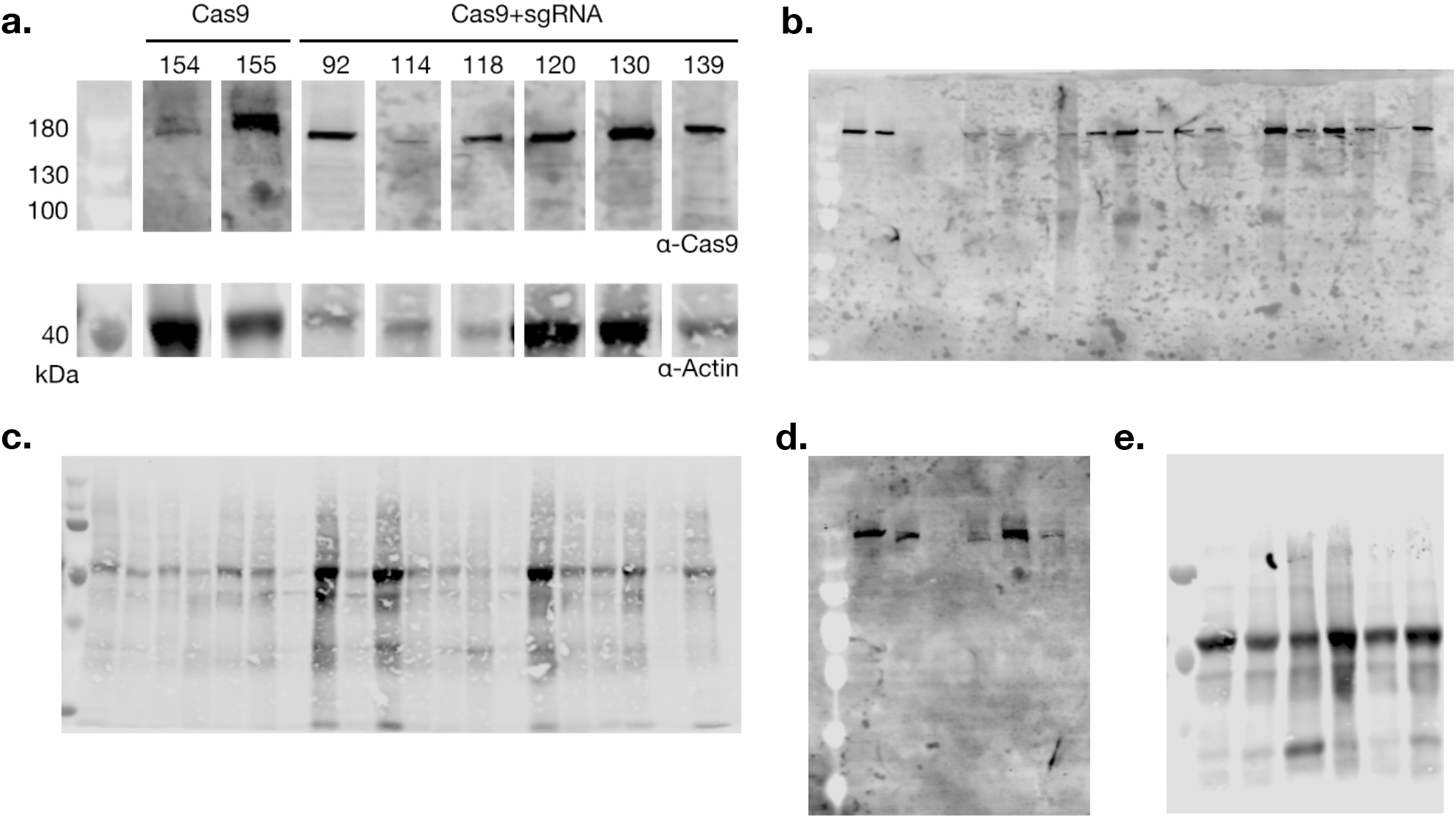
**(a)** Western blots for Cas9-GFP expression. **(b-e)** Raw blot images acquired using an Odyssey CLX imager for anti-Cas9 and anti-Actin antibodies for Cas9+sgRNA1 and Cas9 lines respectively.

**Supplementary Figure 9:**
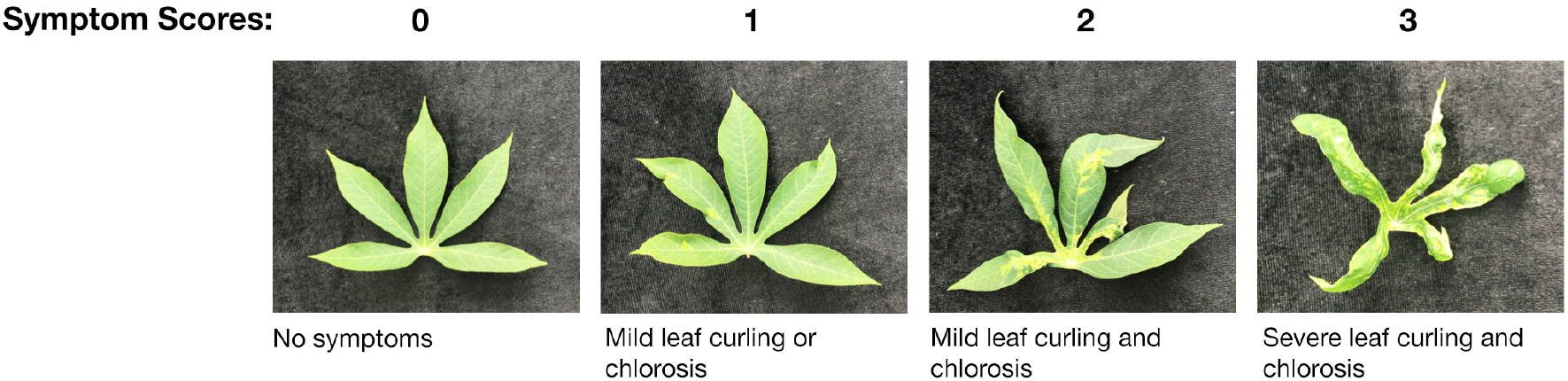
Symptom scoring scale.

**Supplementary Figure 10:**
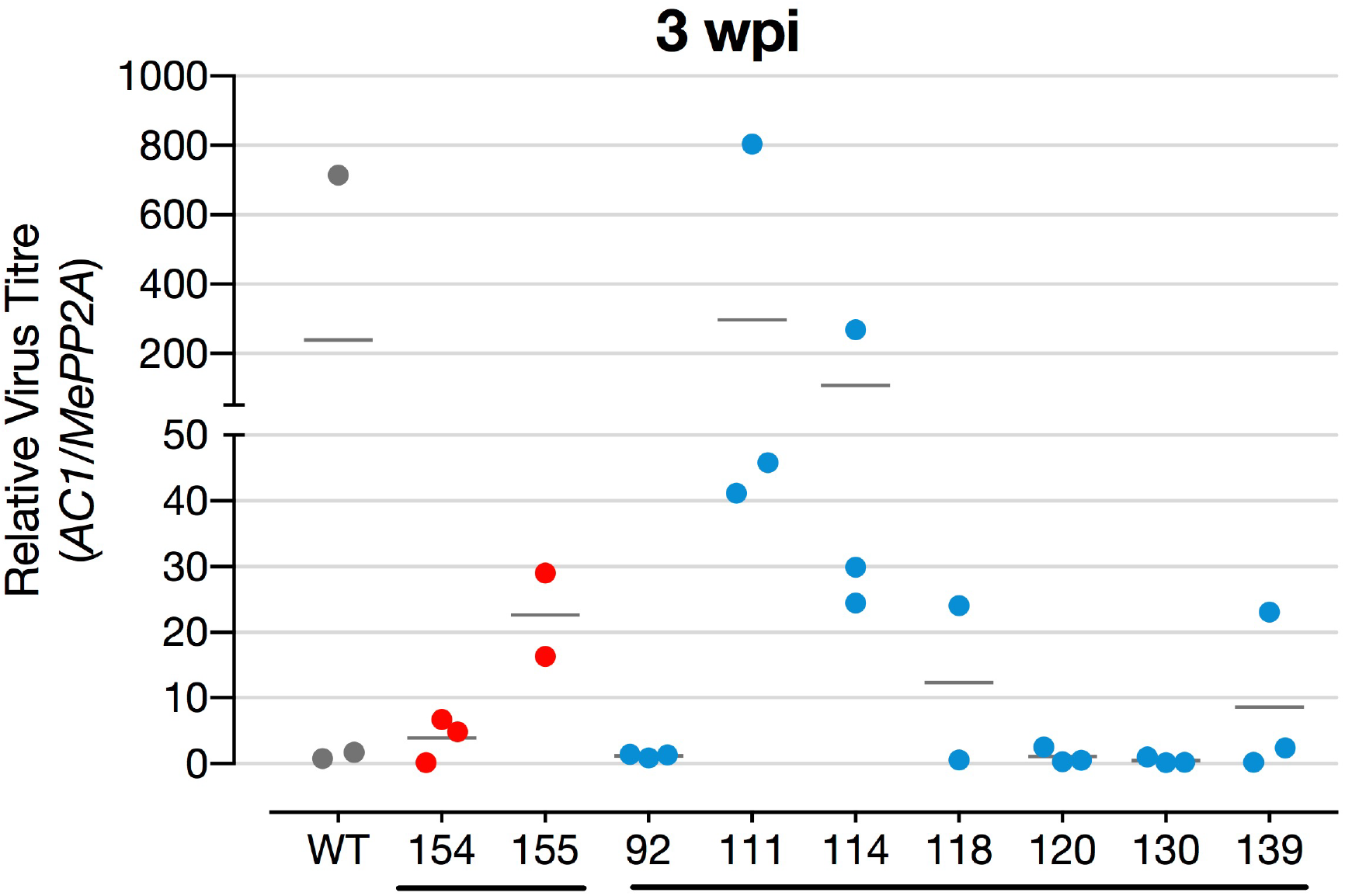
Virus levels in symptomatic plants at 3 weeks post infection measured by qPCR. Levels were measured on replicated pools (each data point in the graph) of leaf samples from a total of at least 4 plants per line.

**Supplementary Figure 11:**
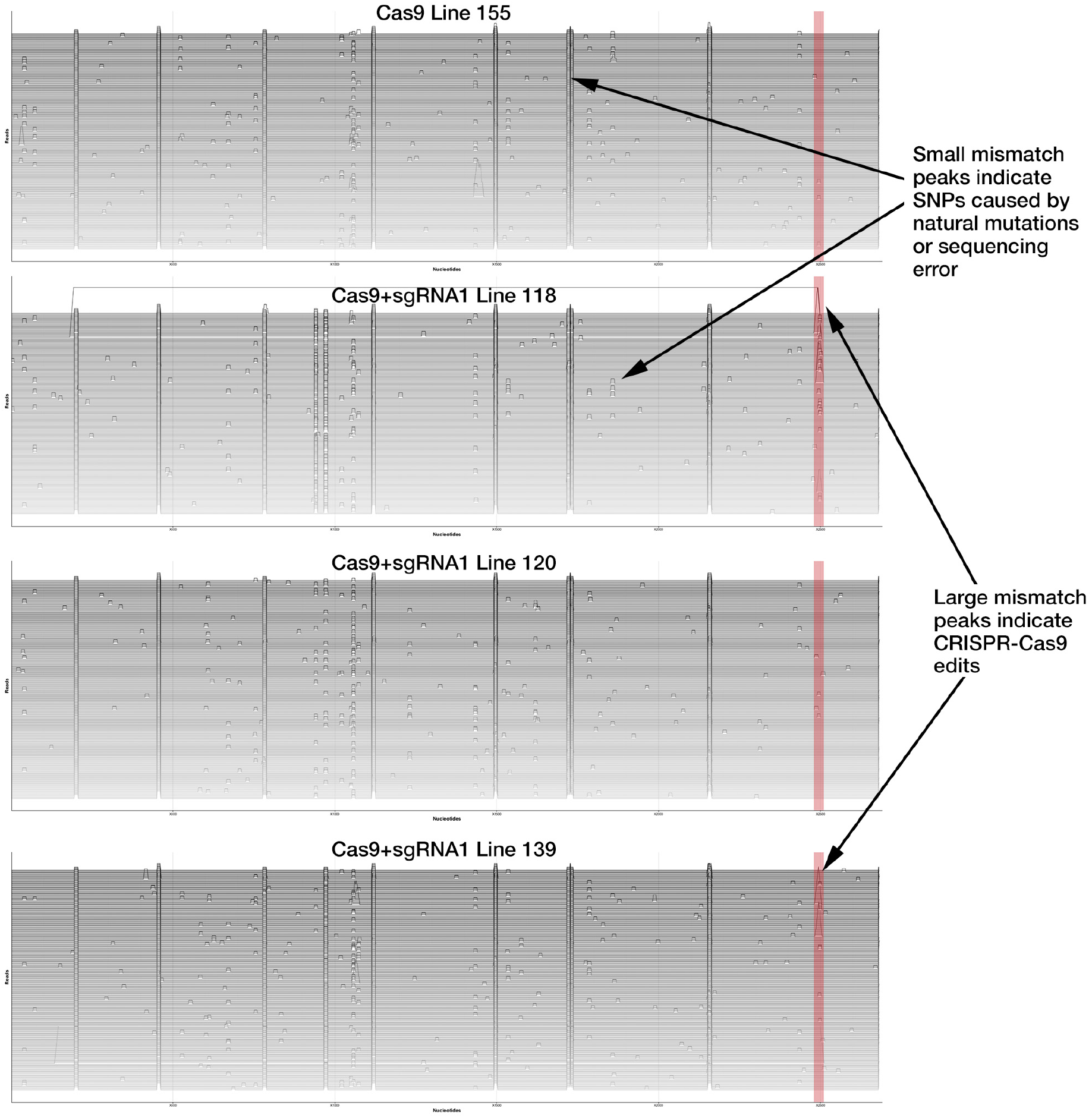
Analysis of full-length virus sequences from infected plants at 8 weeks post infection. Each horizontal line represents a near full-length individual virus sequence (2690bp). Peaks represent edits and are scaled to the %mismatch value of each base-pair (see Methods for calculation) in a pairwise global alignment with the reference virus sequence. The sgRNA target is indicated by a shaded red rectangle. The absence of CRISPR-Cas9 edits (large, tall peaks) in non-target regions indicates a lack of significant off-target cleavage.

### Supplementary Tables

**Supplementary Table 1:**
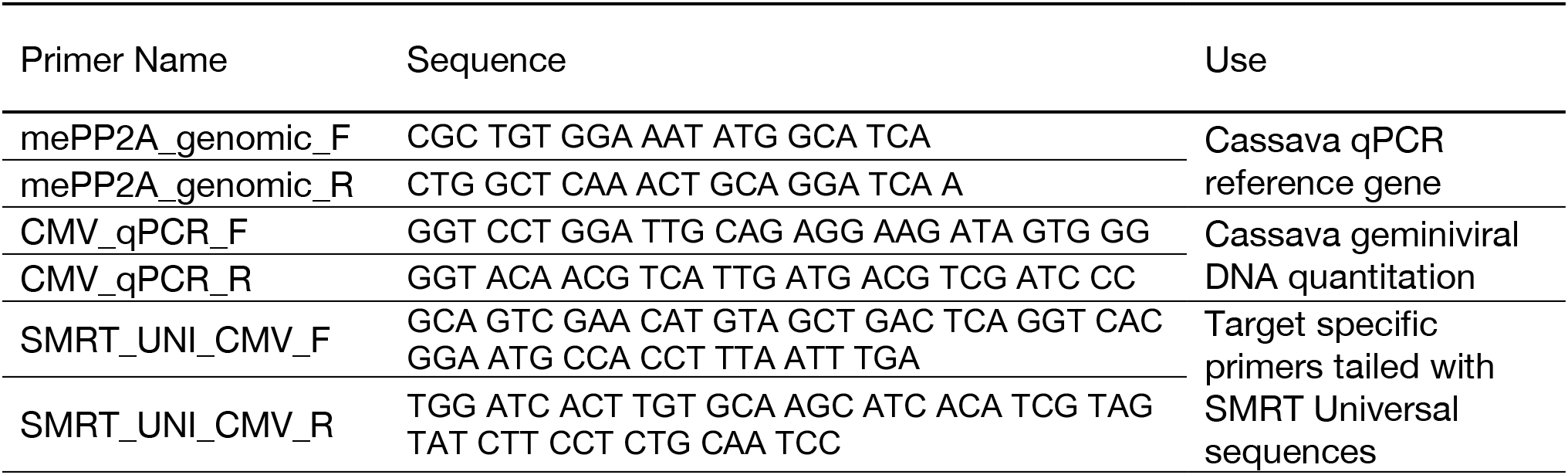
Primer sequences

**Supplementary Table 2:** Sequencing Barcodes

| Barcode | Line | Time-point<br>(weeks after infection) |
| --- | --- | --- |
| 1 | WT | 3 |
| 9 | 155 | 3 |
| 49 | 139 | 3 |
| 41 | 130 | 3 |
| 33 | 118 | 3 |
| 25 | 114 | 3 |
| 17 | 111 | 3 |
| 57 | WT | 8 |
| 73 | 92 | 8 |
| 65 | 155 | 8 |
| 34 | 139 | 8 |
| 26 | 130 | 8 |
| 18 | 120 | 8 |
| 10 | 118 | 8 |
| 2 | 114 | 8 |
| 89 | 111 | 8 |
| 81 | 110 | 8 |

**Supplementary Data 1.**
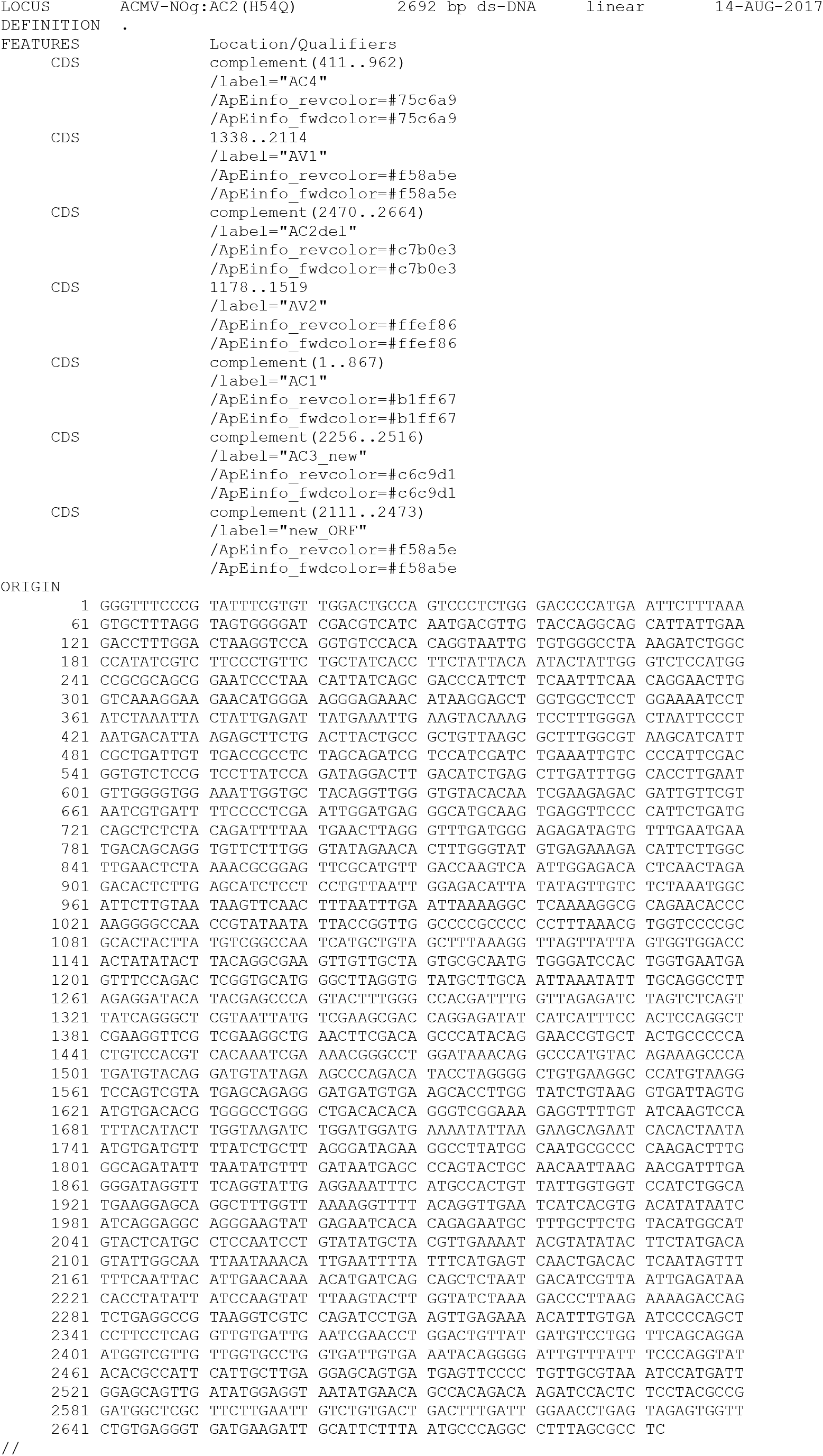

**Supplementary Data 2.**
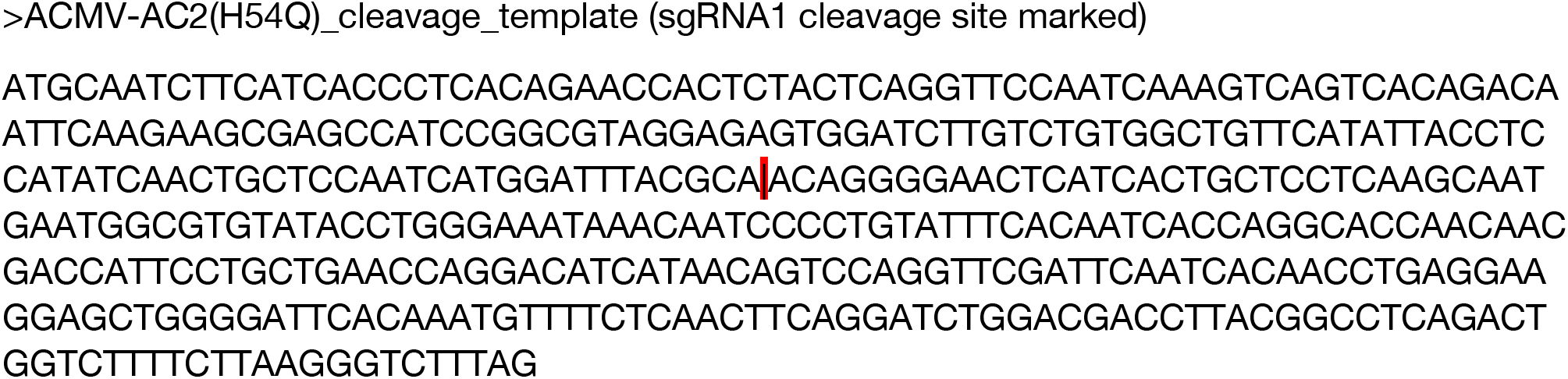

## REFERENCES

1. Kennedy, E. M. & Cullen, B. R. Bacterial CRISPR/Cas DNA endonucleases: A revolutionary technology that could dramatically impact viral research and treatment. Virology 479–480, 213–220 (2015).

2. Baltes, N. J. et al. Conferring resistance to geminiviruses with the CRISPR–Cas prokaryotic immune system. Nat. Plants 1, 15145 (2015).

3. Ji, X., Zhang, H., Zhang, Y., Wang, Y. & Gao, C. Establishing a CRISPR–Cas-like immune system conferring DNA virus resistance in plants. Nat. Plants 1, 15144 (2015).

4. Ali, Z. et al. CRISPR/Cas9-mediated viral interference in plants. Genome Biol. 16, 238 (2015).

5. Chaparro-Garcia, A., Kamoun, S. & Nekrasov, V. Boosting plant immunity with CRISPR/Cas. Genome Biol. 16, 254 (2015).

6. Rey, C. & Vanderschuren, H. Cassava Mosaic and Brown Streak Diseases: Current Perspectives and Beyond. Annu. Rev. Virol. 4, 8.1-8.24 (2017).

7. Vanderschuren, H., Alder, A., Zhang, P. & Gruissem, W. Dose-dependent RNAi-mediated geminivirus resistance in the tropical root crop cassava. Plant Mol. Biol. 70, 265–272 (2009).

8. Burgyán, J. & Havelda, Z. Viral suppressors of RNA silencing. Trends Plant Sci. 16, 265–72 (2011).

9. Doench, J. G. et al. Rational design of highly active sgRNAs for CRISPR-Cas9–mediated gene inactivation. Nat. Biotechnol. 32, 1262–7 (2014).

10. Hanley-Bowdoin, L., Bejarano, E. R., Robertson, D. & Mansoor, S. Geminiviruses: masters at redirecting and reprogramming plant processes. Nat. Rev. Microbiol. 11, 777–788 (2013).

11. Shuman, S. & Glickman, M. S. Bacterial DNA repair by non-homologous end joining. Nat. Rev. Microbiol. 5, 852–861 (2007).

12. Bikard, D. et al. Exploiting CRISPR-Cas nucleases to produce sequence-specific antimicrobials. Nat. Biotechnol. 32, 1146–50 (2014).

13. Citorik, R. J., Mimee, M. & Lu, T. K. Sequence-specific antimicrobials using efficiently delivered RNA-guided nucleases. Nat. Biotechnol. 32, 1141–1145 (2014).

14. Kennedy, E. M. et al. Suppression of hepatitis B virus DNA accumulation in chronically infected cells using a bacterial CRISPR/Cas RNA-guided DNA endonuclease. Virology 476C, 196–205 (2014).

15. Ebina, H., Misawa, N., Kanemura, Y. & Koyanagi, Y. Harnessing the CRISPR/Cas9 system to disrupt latent HIV-1 provirus. Sci. Rep. 3, 2510 (2013).

16. Zhou, X. et al. Evidence that DNA-A of a geminivirus associated with severe cassava mosaic disease in Uganda has arisen by interspecific recombination. J. Gen. Virol. 2101–2111 (1997).

17. Engler, C. et al. A Golden Gate modular cloning toolbox for plants. ACS Synth. Biol. 3, 839–843 (2014).

18. Anders, C. & Jinek, M. in. Methods in Enzymology 0–21 (2014).

19. Bull, S. E. et al. Agrobacterium-mediated transformation of friable embryogenic calli and regeneration of transgenic cassava. Nat. Protoc. 4, 1845–1854 (2009).

20. Chang, S., Puryear, J. & Cairney, J. A simple and efficient method for isolating RNA from pine trees. Plant Mol. Biol. Report. 11, 113–116 (1993).

